# Elevated mutation in haploid yeast driven by translesion synthesis

**DOI:** 10.64898/2026.01.22.701062

**Authors:** Jacob Fredette-Roman, Denise R. Smith, Sanad B. Omari, Nathaniel P. Sharp

**Affiliations:** University of Wisconsin-Madison, Department of Genetics

## Abstract

The impact of selection versus genetic drift on the evolution of mutation patterns is unclear. In *Saccharomyces cerevisiae*, which is predominantly diploid in nature, there is evidence that haploid cells have a higher mutation rate than diploids, suggesting that a haploid-specific mutator phenotype may have evolved due to the limited opportunity for selection to act on this rare cell type. Mutation in haploids was primarily elevated in late-replicating regions of the genome, implicating error-prone translesion synthesis (TLS) repair. Additional research has demonstrated that removing *REV1*, a gene responsible for initiating TLS, causes a reduction in haploid mutation rate. To assess whether the preferential use of this error-prone repair pathway by haploids explains the difference in genome-wide mutation patterns between cell types, we deleted *REV1* in both diploid and haploid *S. cerevisiae* and estimated their mutation rates using a mutation accumulation experiment. Consistent with a previous study, we found a 50% higher single nucleotide mutation rate in *REV1*+ haploids than in *REV1*+ diploids. Deleting the *REV1* gene caused this difference to vanish, with mutation rates in haploid and diploid *rev1*Δ lines converging on 2.4 × 10^−10^. Our results suggest that the mutagenic effect of translesion synthesis is much stronger in haploids, reflecting a limited opportunity for selection to act on mutation rates in rarer cells or smaller populations. We also find evidence that *REV1* plays an important role in mitochondrial genome maintenance in both cell types.

## Introduction

Spontaneous mutations provide the fuel for evolutionary change but can also erode the health of individuals and populations. While the evolutionary significance of mutations is well appreciated, many questions remain about the factors influencing the evolution of mutation rates and spectra. Comparisons of single-nucleotide mutation (SNM) rates among species have revealed that they vary by orders of magnitude across the tree of life (Lynch et al., 2023; Wang & Obbard, 2023). One explanation for this large-scale variation is the drift-barrier hypothesis, which posits that purifying selection generally acts to reduce mutation rates but is unable to prevent the spread of mutator alleles whose fitness effects are weak relative to the effective population size (Lynch et al., 2016). While the drift-barrier hypothesis helps explain why mutation rates vary among species, it remains a challenge to identify the specific genetic factors involved. Importantly, the drift barrier hypothesis could also apply within a species if some DNA replication and repair pathways are rarely deployed and therefore subject to weaker effective selection. There is evidence for such an effect in the budding yeast *Saccharomyces cerevisiae*.

*S. cerevisiae* can be cultured in the lab as a haploid or a diploid, but wild strains are generally diploid (Peter et al., 2018), sporulating to form haploid cells briefly when nutrients become limited (Herskowitz, 1988; Hittinger, 2013). Laboratory strains remain haploid only as long as mating type switching is prevented by genetic manipulation and they are grown without mating partners. While the use of haploid strains is often advantageous for genetic studies, haploid and diploid cells are not necessarily identical, and ploidy-specific variation has been detected for gene expression (Galitski et al., 1999), stress tolerance (Anderson et al., 2004; B.-Z. Li et al., 2010; Mercier, 2005) and DNA repair patterns (Skoneczna et al., 2015). Given that DNA replication in this organism has occurred in the haploid state much less often than the diploid state over evolutionary history, alleles causing elevated mutation in haploids would be largely hidden from selection, allowing the haploid mutation rate to increase due to genetic drift.

Consistent with these ideas, Sharp et al. (2018) found that haploid *S. cerevisiae* accumulated SNMs at a rate 40% higher than diploids from the same genetic background. This pattern also holds in the fission yeast *Schizosaccharomyces pombe*, in which diploids, which are the rarer cell type, show a 60% higher mutation rate than haploids (Bao et al., 2025). In the case of *S. cerevisiae*, the ploidy difference in mutation was strongly driven by genome replication timing: while the SNM rate did not differ between haploids and diploids in the earliest replicating parts of the genome, later-replicating regions showed a significant difference in mutation rate and spectrum (Sharp et al., 2018). This pattern suggests that haploids and diploids differentially regulate DNA repair activities occurring later in the cell cycle, leading us to investigate the role of *REV1*, which plays a role in translesion synthesis (TLS) repair.

When DNA lesions are encountered near the end of the replication process, cells can employ TLS polymerases, including the Rev1 polymerase complex, to replicate across damaged or distorted templates in a time-efficient but error-prone solution to stalled replication (Lehmann, 2006; Sale, 2013; Smith et al., 2007). Rev1 localizes to both the nucleus and the mitochondria and has been found to be under cell-cycle control in haploids, with elevated levels in the G_2_ phase (Waters & Walker, 2006; Zhang et al., 2006). A study using haploid *S. cerevisiae* found that mutation rates in *URA3* were elevated when the gene was inserted into later-replicating regions of the genome, but that this effect was suppressed by the deletion of *REV1* (Lang & Murray, 2011), meaning that *REV1*-initiated TLS was responsible for an elevated mutation rate in late-replicating regions of the genome in haploids. Combined with the findings from Sharp et al. (2018), we therefore hypothesized that greater use of this error-prone repair pathway in haploids could account for the higher mutation rate in this cell type overall.

We predicted that deleting *REV1* in both haploids and diploids would eliminate the ploidy difference in mutation rate. To test this, we conducted mutation accumulation (MA) on lines of *REV1*+ and *rev1*Δ haploid and diploid *S. cerevisiae* derived from a common genetic background. We maintained 200 lines under serial single-cell bottlenecks for a total of ∼141,000 generations. This approach limited the effective population size (N_e_) in our lines, such that most new mutations would have approximately the same probability of fixation as a neutral allele (Halligan & Keightley, 2009), allowing us to measure mutation patterns without bias due to selection.

Whole genome sequencing of our MA lines revealed clear impacts of ploidy and *REV1* genotype on mutation patterns in both the nuclear and mitochondrial genomes. Our results show that deleting a single DNA repair gene in *S. cerevisiae* can cause a ploidy difference in SNM rates to disappear, suggesting that TLS use represents a haploid-specific mutator phenotype. To further explore the role of Rev1 in DNA repair in haploids and diploids, we also examined how the absence of *REV1* impacted resistance to UV exposure and tert-butyl hydroperoxide (t-BOOH), two commonly used mutagens.

## Materials and Methods

### Yeast strains

We obtained the haploid strain SEY6211 (*MAT*a *leu2-3*,*112 ura3-52 his3-*Δ*200 trp1-*Δ*901 ade2-101 suc2-*Δ9 *ho*) from the American Type Culture Collection and used a single-cell isolate to generate our experimental strains. We used a standard plasmid transformation protocol to induce mating type switching, then crossed *MAT*a and *MATα* haploids to generate a diploid strain. To delete *REV1* (positions 981828-984785 on chromosome XV) we transformed haploid *MAT*a cells with a *KanMX* cassette flanked by regions of homology with the relevant genomic region using the lithium acetate method (Gietz & Schiestl, 1991), and selected for integrants on geneticin (G418). We confirmed the deletion of *REV1* using long-read amplicon sequencing. After deleting *REV1* in the *MAT*a haploid, we repeated the mating type switch and crossed *MAT*a and *MATα* cells to generate a *rev1*Δ diploid. We preserved these strains as ancestral controls and isogenized to single colonies to initiate the bottlenecking procedure. We used only diploids and *MAT*a cells for MA, since there is no evidence for a difference in mutation patterns between *MAT*a and *MATα* haploids (Sharp et al. 2018).

### Mutation accumulation

We conducted MA transfers on solid yeast-extract-peptone-dextrose (YPD) media, on 5.5 cm diameter plates. We incubated plates at 30° C and performed single-cell bottlenecks three times weekly for a total of 30 bottlenecks. We retained older plates at 4° C as backups but did not have to use any backup plates.

### DNA extractions

Following MA, we extracted DNA from the MA lines and the ancestral strains using the ZymoResearch YeaStar Genomic DNA Kit with the chloroform protocol (Zymo Research, Irvine, CA) and quantified it using the Qubit dsDNA HS Assay Kit (Life Technologies, Grand Island, NY). Libraries were prepared by the University of Wisconsin-Madison Biotechnology Center using the QIAGEN FX DNA Library Preparation Kit (QIAGEN) and sequenced on an Illumina NovaSeq6000 system with paired-end 150-bp reads (Illumina, San Diego, CA). The average coverage per line before screening was 106X for haploids and 107X for diploids.

### Sequence analysis

We used BBduk v39.01 and Trimmomatic v0.39 to trim adapters and low-quality ends, respectively, and used FastQ Screen v0.15.3 to verify that contamination was absent in our samples. We aligned trimmed reads to the reference genome (assembly R64-1-1) using bwa-mem2 v2.2.1 (H. Li, 2013), removed duplicate reads using Picard MarkDuplicates v3.0.0 (Broad Institute, 2020), and generated gVCF (genomic Variant Call Format) files using HaplotypeCaller, CombineGVCFs, and GenotypeGVCFs in GATK v4.4.0.0 (Poplin et al., 2017). We used GATK SelectVariants to screen single-nucleotide variants (SNV) and indels separately, and applied hard filters to each callset using the GATK best practices workflow for germline short variant discovery (Auwera & O’Connor, 2020). We excluded any variants that failed hard filters or were present in multiple MA lines or the ancestral control strains.

We estimated mutation rates using *μ* = *m*/(*c g*), where *m* is the number of variants in the final callset, *c* is the number of callable sites across all lines for a given treatment (multiplied by two for diploids), and *g* is the number of generations of MA. For a site to be considered callable in a given sample, we required that (i) GATK called a genotype for the sample at that site and (ii) GATK called genotypes for at least 90% of all MA samples at that site.

We classified any SNVs within 100 bp of one another in the same sample as a multi-nucleotide mutation event (MNM), with each cluster counted only once for the purposes of mutation rate estimation. We classified cases where at least one variant in a cluster was an indel as “complex” mutations. Unless otherwise noted, our SNM rate estimates exclude MNMs, and our indel rate estimates include complex events. We determined the genic context and impact of mutations using Ensembl Variant Effect Predictor (McLaren et al., 2016). We detected aneuploidy by comparing median read depth across chromosomes in each sample; if the median depth within a chromosome was >1.3x or <0.7x the average of median depths for all chromosomes in a sample, we called a chromosome gain or loss, respectively.

### Flow cytometry

To assess ploidy following MA, we stained DNA in fixed cells with SYTOX Green and measured fluorescence using an Attune NxT flow cytometer following Sharp et al. (2018). We tested MA lines in a single block of three 96-well plates, along with three replicates of each ancestral control, analyzing approximately 27 000 cells per line. We removed any data points with signal pulse heights above 4 × 10^5^ and below 2.5 × 10^4^ to filter out any potential debris, bacterial cells, or flocculating yeast cells. To determine ploidy, we visually compared fluorescence peaks in MA lines with those of the ancestral controls. In cases where the ploidy of an MA line deviated from its starting ploidy or ploidy was difficult to ascertain from the data, we re-ran samples as a second block with new controls and analyzed the data in the same way.

### Phenotype testing

Following MA, we re-initialized MA lines as patches on YPD plates and replica-plated onto non-fermentable yeast-extract-peptone-glycerol (YPG) medium to evaluate whether any lines had become *petite* mutants. We also replica-plated patches onto media containing G418 to verify the status of *REV1*. Finally, we performed test crosses for each MA line with MATa and MAT*α* tester strains and plated on minimal media to determine mating type.

### Growth rate measurements

Following MA, we determined the number of generations of MA for each set of lines by re-initializing the ancestral strains and five random MA lines from each treatment on YPD plates, streaking to single colony. Following 48 and 72 hours of growth, we combined six random colonies from each plate in sorbitol and measured cell concentrations using a hemocytometer to infer the average number of cells per colony, *n*. We estimated the rate of doubling over *t* hours as *r_t_* = log_2_(*n*)/*t* and estimated the cell division rate for a given treatment as (*r_t,MA_* + *r_t,control_*)/2 for that treatment. We estimated the generations of MA for each treatment by finding the weighted average of the 48- and 72-hour cell division rate estimates at a ratio of 2:1 (reflecting two 48-hour transfers and one 72-hour transfer per week of MA) and multiplying that rate by 70 days of MA.

### Mutagen assays

To assess the relative fitness of our genotypes of interest in the presence of oxidative stress, we competed REV1+ and *rev1*Δ cells in the presence or absence of tert-butyl hydroperoxide (t-BOOH), a commonly used oxidative stressor (Kučera et al., 2014). For both haploids and diploids, we combined REV1+ and *rev1*Δ cells in replicate liquid cultures, beginning with 10 000 cells of each type in either 2 mL of YPD or YPD with 1 mM t-BOOH. Following 24 hours of growth at 30° C, we spread cells from these cultures on YPD plates, allowed colony growth, and then patched random colonies onto media containing G418. Since the *rev1*Δ strains are G418 resistant, we could estimate their frequency as the proportion of patches that grew on the G418 plates.

We also assessed the survival of each genotype of interest following exposure to ultraviolet radiation. Following 48 hours of growth in liquid YPD, we diluted cultures to a concentration of 1 cell/μL and plated 200 μL on replicate YPD plates. We exposed half of the plates to irradiation (155 µW/cm^2^ UVA, 56 µW/cm^2^ UVB and 36 µW/cm^2^ UVC) for 2 minutes, with the other half serving as controls. Following 48 hours of growth at 30° C, we counted colonies on each plate and estimated UV resistance as the number of colonies on the UV-treated plates divided by the number of colonies on the control plates.

### Statistical comparisons

We conducted statistical comparisons using *R* 4.5.1 (R Core Team, 2025). In the case of significant interaction effects in generalized linear models, we conducted post-hoc pairwise comparisons using estimated marginal means with the *emmeans* package (Lenth, 2025). To determine the replication timing of variable sites, we used data from Müller et al. (2014).

## Results

We examined the mutation profiles of haploids and diploids with and without a functional copy of the *REV1* gene. Following MA, we identified 1,003 nuclear mutations and 79 mitochondrial mutation events (Figure 1). Within the nuclear genome, we found patterns associated with specific types of point mutations (Figure 2), and we analyzed the effect of ploidy and *REV1* status on growth rate and tolerance to exogenous mutagens (Figure 3).

**Figure 1.**
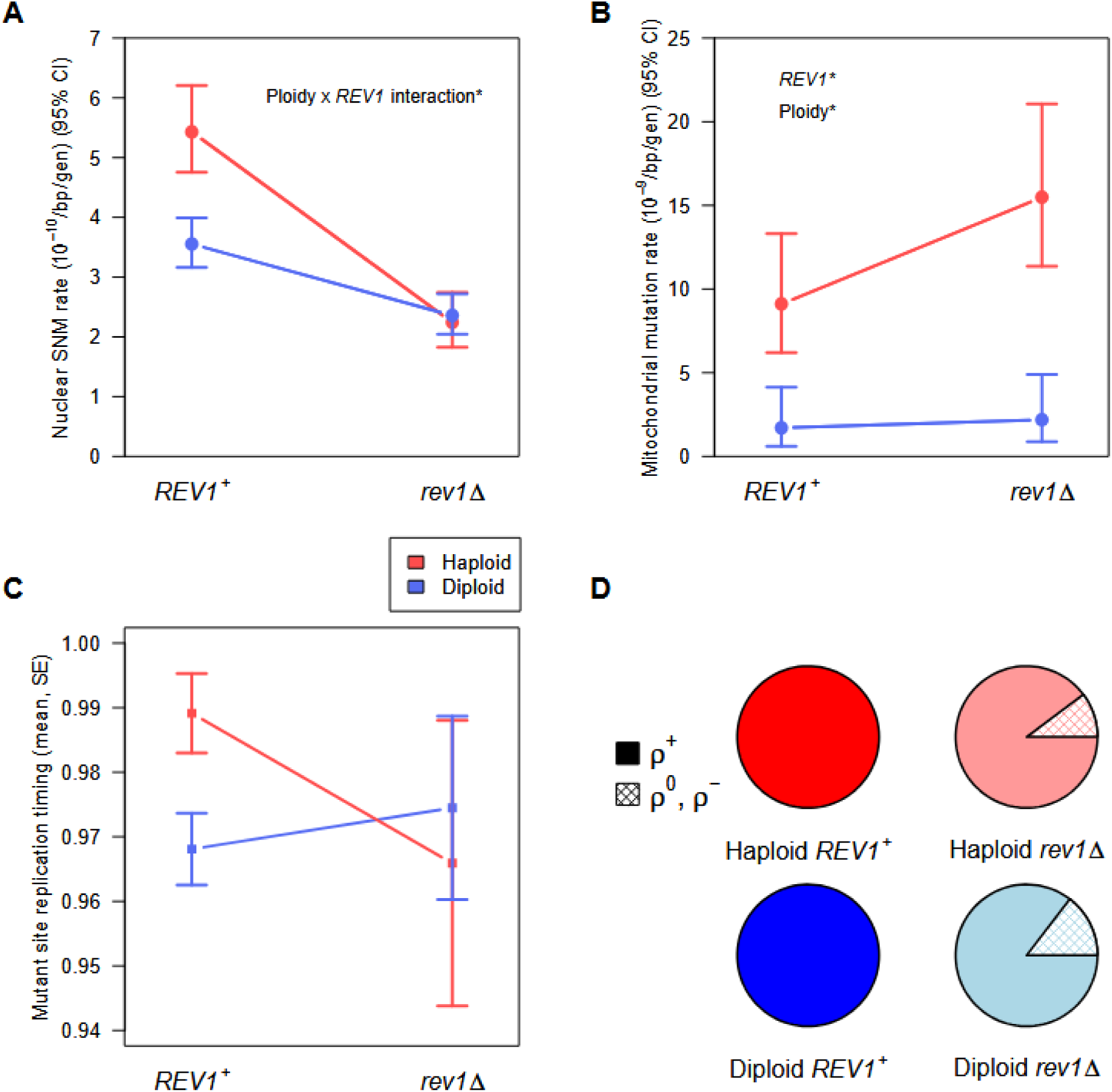
Nuclear and mitochondrial mutation rate estimates, and mean replication timing of nuclear variants called. (A) Among wild-type lines, haploid yeast had a higher single-nucleotide mutation (SNM) rate compared to diploids. Deleting *REV1* caused SNM rates to decrease significantly in both haploids and diploids. We detected a significant interaction effect between ploidy and *REV1* status on SNM rate, where *REV1* deletion had a larger effect on haploids. (B) The rate of mutations called in the mitochondrial genome was overall higher in haploids and in *rev1*Δ treatments, but we did not detect an interaction effect between the two factors. (C) Mean replication timing of variants called across all four treatments, with the *REV1+* group including variants called in Sharp et al. (2018). We did not detect significant effects treatment on the mean replication timing of mutations, likely due to the limited sample size of variants called in *rev1*Δ treatments. (D) Deleting the *REV1* gene significantly increased the rate that lines became petite in both haploid and diploid treatments. None of the 100 total *REV1+* lines became petite over the course of MA. We calculated 95% binomial confidence intervals using the Clopper-Pearson exact method, and we excluded lines that changed ploidy.

**Figure 2.**
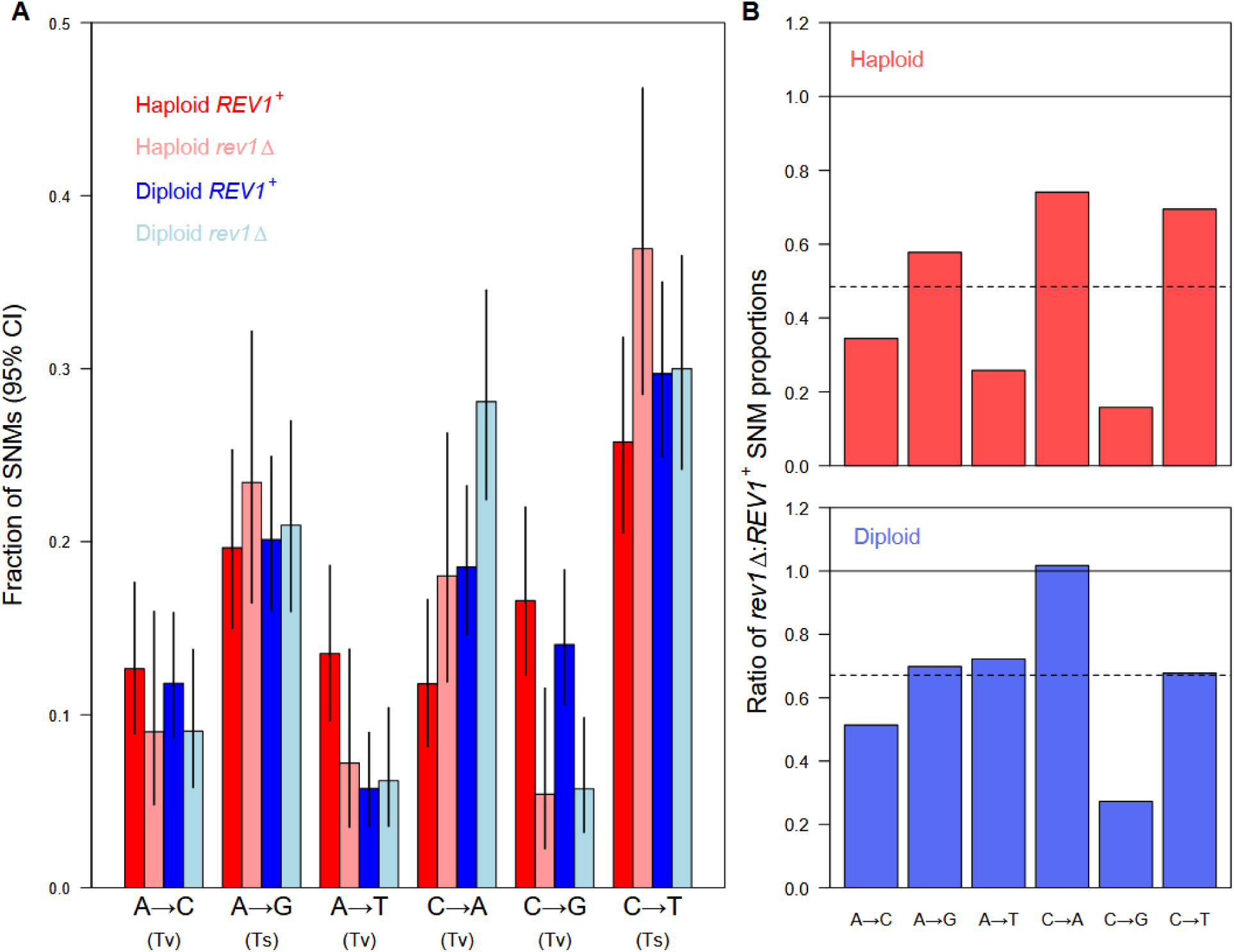
Point mutation spectrum in yeast grown as haploids or diploids, with *REV1* present or deleted. (A) The proportion of mutations across six categories differed significantly between the four treatment groups. Specifically, both haploids and diploids showed a significantly reduced share of C to G transversions following the deletion of *REV1*. 95% binomial confidence intervals were calculated using the Clopper-Pearson exact method. (B) The ratio of SNM proportions from (A) between *rev1*Δ and *REV1+* treatments in haploids (top) and diploids (bottom). Solid lines are shown at a ratio of 1, which would be expected if SNM counts between *rev1*Δ and *REV1+* groups did not differ within a given ploidy level. Dotted lines are shown at ratios that reflect the expectation if each SNM type differed between *rev1*Δ and *REV1+* groups equally within a given ploidy. We excluded SNMs belonging to lines that changed ploidy for this analysis.

**Figure 3.**
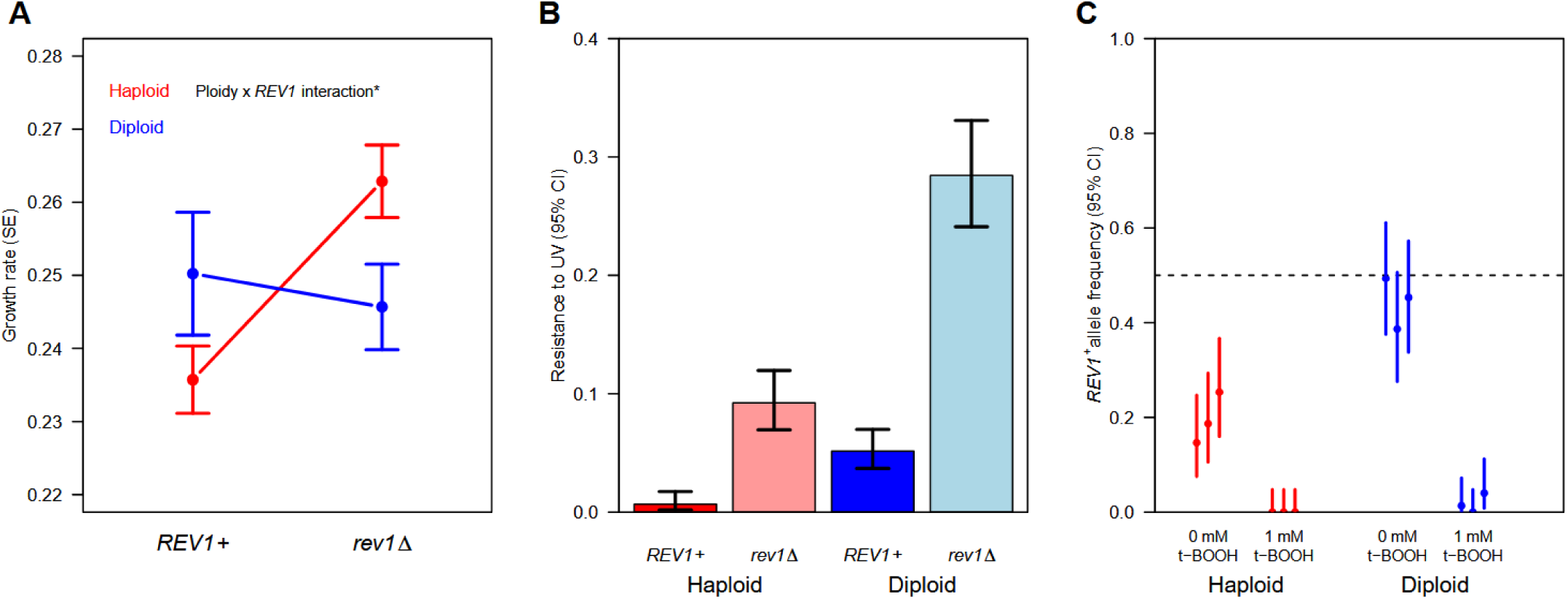
Growth of haploid and diploid *REV1*+ and *rev1*Δ strains with and without exposure to mutagenic stressors. (A) Deleting *REV1* caused an increase in growth rate among haploid cells growing in nutrient-rich liquid media but had no effect in diploids. The growth rate of *rev1*Δ haploids was statistically indistinguishable from the growth rate of *REV1* diploids. (B) Deleting the *REV1* gene caused resistance to UV to increase in both haploids and diploids. We also detected an interaction effect between ploidy and *REV1* genotype on UV resistance. (C) *rev1*Δ strains outcompeted *REV1+* strains in both ploidy types, and this effect was magnified upon the introduction of 1mM t-BOOH. We found evidence for main effects of both t-BOOH and ploidy on the relative proportion of surviving *REV1*+ cells following 24 hours of competitive growth.

### Ploidy and phenotype testing

Following MA, all 200 lines retained the G418 resistance status of the ancestral strain from which they originated, but we found evidence for ploidy change in two cases. Specifically, two initially-haploid MA lines (one *REV1*+ and one *rev1*Δ) showed DNA levels consistent with diploidy based on flow cytometry (Table S1, Fig. S1). These lines showed *MAT*a-like mating behavior, indicating that they most likely became *MAT*a/*MAT*a diploids through endoreduplication (Gerstein & Sharp, 2021; Harari et al., 2018). Our data indicate a diploidization rate of 1.24 × 10^−5^ per haploid cell division (95% CI: 0.149 – 4.47 × 10^−5^). We excluded these two lines from further analysis. We also found that two lines from the diploid *rev1*Δ group showed haploid-like mating behavior in test crosses (one *MAT*a-like and one *MAT*⍺-like). These lines remained diploid in terms of DNA content, with no evidence of aneuploidy, so the most likely explanation is loss of heterozygosity (LOH) at the *MAT* locus. Our data indicate that LOH at *MAT* occurs at a rate of 1.25 × 10^−5^ per diploid cell division (95% CI: 1.51 × 10^−6^ – 4.52 × 10^−5^), consistent with previous reports (Harari et al., 2018). Given that gene expression patterns can vary according to the state of the *MAT* locus (Anderson et al., 2004; Galitski et al., 1999; B.-Z. Li et al., 2010), we also excluded these lines from further analysis.

We also found that 5 haploid lines and 7 diploid lines became petite and did not grow on YPG, a non-fermentable carbon source, following MA (Fig. 1D). Notably, all 12 of these lines belonged to *rev1*Δ treatments. Ploidy did not affect the rate of petiteness over the course of MA (binomial GLM, *z* = –0.65, *P* = 0.51), but lines from the *rev1*Δ treatments were overall more likely to be petite (Fisher’s Exact Test, *P =* 1.5 × 10 ^−4^). Petiteness did not significantly affect estimates of single-nucleotide mutation rate across *rev1*Δ treatments (binomial test, *P* = 0.11), so petite lines were included in nuclear and mitochondrial mutation rate comparisons. However, most of the petite lines lacked mitochondrial genomes entirely (see below).

We observed a pink pigmentation in colonies formed by both haploid and diploid *REV1*+ ancestral strains, consistent with previous studies on the SEY6211 strain (Wang et al., 2001), but this phenotype was absent in *rev1*Δ strains. Among our ancestral sequences belonging to these strains, we observed a C>A change on position 865,708 of chromosome XIII that was not present in the *REV1*+ background. According to Variant Effect Predictor, this mutation corresponds to a p.E462* change in *ADE4*. This could have contributed to the pigmentation change, as disruption to the purine nucleotide biosynthetic pathway caused by *ADE2* mutants have been shown to cause an accumulation of red pigment in cells deprived of adenine (consistent with the original SEY background), while cells with nonfunctional *ADE2* and *ADE4* genes do not accumulate a red pigment (Nevzglyadova et al., 2022). We are not aware of any evidence that this pigment would influence genome maintenance in rich media.

### Growth rates

We found a significant decrease in growth rate in MA lines relative to ancestral strains regardless of ploidy (381 colonies measured; linear model, *t* = 2.92, *P* = 0.004). While we did not detect a growth rate difference between haploid and diploid treatments (*t* = 1.168, P = 0.24), we observed a ∼4% higher growth rate in *rev1*Δ lines in both haploids and diploids grown on YPD (*t* = –4.69, *P* < 10^−6^). Growth rate data for MA lines and ancestral controls are provided in Table S2. To assess the possibility that this growth rate difference could have been caused by the loss of function mutation in the *ADE4* gene rather than the deletion of *REV1*, we measured maximum rates of population growth in liquid YPD supplemented with adenine (YPAD) using a Biotek Epoch II plate reader. With supplemental adenine, mutations in the de novo purine nucleotide biosynthetic pathway should be less relevant (Ljungdahl & Daignan-Fornier, 2012; Som et al., 2005). We detected a significant interaction between ploidy and *REV1* genotype in this assay (Fig. 3A; linear model, *t* = –2.58, *P* = 0.011), with the deletion of *REV1* associated with faster growth in haploids (*t* = –4.01, *P* = 2.23 × 10^−4^) and no change in diploids (*t* = 0.44, *P* = 0.66). This means that while the presence of *REV1* reduced growth rate in haploids, we observed no such effect in diploids. The growth rate of *rev1*Δ haploids was statistically indistinguishable from that of *REV1* diploids (*t* = 1.29, *P* = 0.20).

### Nuclear mutations

We called 1,003 total mutations in the nuclear genome and classified them as single-nucleotide mutations (SNMs), indels, multi-nucleotide mutations (MNMs), or complex mutations based on their proximity to other variants. We detected a significant interaction effect between ploidy and *REV1* status on SNM rate (binomial generalized linear model: *z* = 3.06, *P =* 0.002); the deletion of *REV1* reduced the SNM rate in haploids more than in diploids. This is what we expected to observe if *REV1* was responsible for the difference in haploid and diploid SNM rates observed in Sharp et al. 2018 (Fig. 1A). Post-hoc pairwise comparisons revealed that SNM rates differed significantly between haploids and diploids among *REV1+* treatments (*z* = –4.71, *P* < 0.0001), but not among *rev1*Δ treatments (*P* = 0.69). We estimated the per nucleotide, per generation SNM rate to be ∼50% higher in *REV1+* haploids (5.43 × 10^−10^, 95% CI: 4.75 – 6.20 × 10^−10^) compared to *REV1*+ diploids (3.55 × 10^−10^, 95% CI: 3.16 – 3.99 × 10^−10^), reproducing the main result from Sharp et al. (2018). Upon deleting *REV1*, we observed a 58.7% decline in the SNM rate of haploids (2.24 × 10^−10^, 95% CI: 1.83 – 2.75 × 10^−10^) and a 33.5% decline in the SNM rate of diploids (2.36 × 10^−10^, 95% CI: 2.04 – 2.72 × 10^−10^) (*z* = –7.14, *P* < 0.0001 and *z* = 4.38, *P* < 0.0001, respectively). While we did call several indels, MNMs, and complex mutation events (Table 1), we did not detect any significant differences in the rates of these mutations between our treatment groups. We also identified six aneuploidies across our MA lines, all of which were present in diploids (Table S1). While five of these occurred in *rev1*Δ lines, this rate did not differ significantly from the rate of aneuploidies in the *REV1*+ lines (binomial test, *P* = 0.22).

**Table 1.** Sample sizes, generations, and mutations called.

|  | Haploid |  | Diploid |  |
| --- | --- | --- | --- | --- |
|  | *REV1+* | *rev1Δ* | *REV1+* | *rev1Δ* |
| MA lines | 49 | 49 | 50 | 48 |
| Cell divisions per transfer | 23.2 | 24.1 | 22.9 | 23.9 |
| Transfers per line | 30 | 30 | 30 | 30 |
| Median coverage per site per line | 107 | 106 | 106 | 107 |
| Total callable nuclear sites | 574,165,046 | 574,063,763 | 1,171,426,300 | 1,123,916,736 |
| Single nucleotide mutations | 217 | 93 | 286 | 190 |
| Indels | 20 | 9 | 32 | 31 |
| Multi-nucleotide mutations | 4 | 5 | 6 | 7 |
| Complex mutations | 4 | 3 | 8 | 3 |
| Total callable mitochondrial sites | 4,260,209 | 3,662,262 | 4,259,701 | 3,833,734 |
| Mitochondrial mutation events | 27 | 41 | 5 | 6 |
| Petite lines following MA | 0 | 5 | 0 | 7 |

\*Excludes lines that changed ploidy

### Mitochondrial mutations

Among the 12 MA lines that lost mitochondrial function and became petite, 11 lost most or all of their mtDNA, with median read depths of 0 in the mitochondrial genome alignments. One line, which became petite but retained its mtDNA, acquired an A>T SNM on position 18,264 of the mitochondrial genome. According to Variant Effect Predictor, this mutation corresponds to a p.K666* change in *AI2,* a reverse transcriptase required for the splicing of the *COX1* pre-mRNA encoded within a *COX1* intron.

To further assess the role that *REV1* deletion had on the mitochondrial mutation rate, we examined mitochondrial mutations in lines that did not become petite over the course of MA (Fig. 1B). We did not find a significant interaction effect between ploidy and *REV1* status on the mitochondrial mutation rate (binomial GLM, *z* = –0.44, *P* = 0.66). However, we did detect significant main effects of both ploidy (*z* = 5.66, *P <* 10^−7^) and *REV1* genotype (*z* = – 2.14, *P* = 0.03) on the mitochondrial mutation rate. Among the *REV1*+ treatments, haploids had a mitochondrial mutation rate ∼5 times higher than in diploids (haploid mitochondrial mutation rate: 9.1 × 10^−9^, 95% CI: 6.20 – 13.3 × 10^−9^; diploid mitochondrial mutation rate: 1.71 × 10^−9^, 95% CI: 0.60 – 4.13 × 10^−9^), consistent with previous findings (Sharp et al., 2018; E. A. Sia et al., 2000; R. A. L. Sia et al., 2003). Upon deleting *REV1*, the mitochondrial mutation rate increased 1.7-fold in haploids (1.55 × 10^−8^, 95% CI: 1.14 – 2.11 × 10^−8^) and ∼1.3-fold in diploids (2.18 × 10^−9^, 95% CI: 0.88 – 4.89 × 10^−9^).

### Replication timing

We would expect the deletion of *REV1* to eliminate the mutation bias towards late-replicating regions in haploids (Lang and Murray 2011; Sharp et al. 2018). We found that the distribution of replication timing at mutant sites did not differ between our callset for *REV1*+ strains and the callset from Sharp et al. (2018) for haploid variants (Wilcoxon test, *P* = 0.45) or diploid variants (Wilcoxon test, *P* = 0.23). Therefore, we combined the two datasets for further analyses of replication timing. While the mean replication timing at mutant sites trended downward in haploids and upward in diploids in the *rev1*Δ lines (Fig. 1C), we did not detect a significant interaction effect (randomization test, *P* = 0.24); the lower mutation rate in *rev1*Δ strains likely reduced our power to detect changes in the pattern of mutations with respect to replication timing.

### Mutation spectrum

We detected significant differences between our four treatment groups in the spectrum of SNMs across six categories (Fig. 2A; 4 x 6 chi-square test: χ^2^ = 49.17, *P* < 10^−4^). To determine whether there were any SNM types that differed significantly between *REV1*+ and *rev1*Δ groups within each ploidy level, we first conducted additional 2 x 6 chi-square tests, which indicated significance for both haploids (χ^2^ = 16.49, *P* = 0.0056) and diploids (χ^2^ = 14.30, *P* = 0.014). We examined deviations from expected values (standardized residuals) for each cell in our contingency tables to determine which SNM types were most contributing to the overall difference in mutation patterns. Values less than –2 or greater than 2 indicate significant deviations from expectations (Legendre & Legendre, 2012). Analyses of each 2 x 6 chi-square test revealed that the only significant standardized residuals were in C to G SNMs in *rev1*Δ haploids (standardized residual = –2.21) and C to G SNMs in *rev1*Δ diploids (standardized residual = –2.21). This indicates that *REV1* deletion affected the relative rate of C to G transversions in both haploids and diploids.

We categorized each variant as genic or intergenic, based on the results from Variant Effect Predictor (McLaren et al., 2016) and found no significant difference in the proportion of genic versus intergenic variants between treatments (2 x 4 chi-square test, χ^2^ = 7.07, *P* = 0.07). Among genic variants, we also compared the proportion of nonsynonymous variants and found no significant difference between treatments (χ^2^ = 3.15, *P =* 0.37). We also found no difference between groups in the proportion of genic variants that created nonsense mutations (χ^2^ = 1.2, *P* = 0.75). Taken together, these results suggest that selection did not act differently across treatments.

We observed 8 homozygous SNMs in *rev1*Δ diploids and 14 homozygous SNMs in *REV1*+ diploids, indicating loss of heterozygosity (LOH) events. The proportion of LOH events did not differ by *REV1* treatment (Fisher’s exact test, *P* = 0.83), nor did our overall rate of LOH events differ from the LOH rate reported in Sharp et al. (2018) (2 x 2 chi-square test, χ^2^ = 0.59, *P =* 0.44).

### Mutagen assays

We found that survival on YPD plates was significantly reduced by two minutes of exposure to UV light (Poisson GLM, *z* = –33.2, *P* < 10^−16^). On average, resistance to UV was higher in diploids and higher in *rev1*Δ genotypes; we also detected a significant interaction effect between ploidy and *REV1* genotype (z = –5.44, *P* < 10^−8^), where the *REV1* deletion increased resistance to a greater degree in diploids than in haploids (Fig. 3B).

We also grew our strains competitively in liquid media with and without t-BOOH to test whether *REV1* might offer a competitive advantage under oxidative stress. Following competition in the absence of t-BOOH, both haploid and diploid *rev1*Δ strains outperformed their *REV1*+ counterparts (Fig. 3C), consistent with separate calculations of growth rate used to estimate generations of MA for each group. However, in diploids competing without t-BOOH, the frequency of *rev1*Δ was not significantly different from 0.5 (binomial test, *P* = 0.11), despite diploid *rev1*Δ cells growing faster on solid media in a non-competitive assay (see *Growth rates*). In the presence of 1 mM t-BOOH, the competitive advantage of *rev1*Δ cells became stronger, with zero out of 225 total counted cells across three haploid trials belonging to the *REV1*+ strain, and only four out of 225 cells across three diploid trials belonging to the *REV1*+ strain. While we did not detect a significant interaction effect between ploidy and t-BOOH on the competition assay outcome (binomial GLM, *P* = 0.997), we did find evidence for main effects of both t-BOOH (*z* = –7.26, *P* < 10^−13^) and ploidy (*z* = –5.5, *P* < 10^−8^) on the frequency of *REV1*+ cells.

## Discussion

Across-species comparisons of mutation rate have revealed large-scale patterns of variation corresponding to factors like effective population size, taxonomic domain, and genome size (Conradsen et al., 2022; Lynch et al., 2016; Wang & Obbard, 2023). Additional comparisons of mutation rate in *S. cerevisiae* alone have revealed within-species variation across factors like cell type and environment (Liu & Zhang, 2019; Melde et al., 2022; Sharp et al., 2018), with some studies identifying specific genetic loci that contribute to mutation rate variation across genetic backgrounds (Gou et al., 2019; Jiang et al., 2021). While alleles that contribute to mutation rate variation have been identified in haploid *S. cerevisiae*, this organism is generally diploid in the wild (Bai et al., 2022), and there is evidence that mutation patterns depend on ploidy (Bao et al., 2025; Sharp et al., 2018; E. A. Sia et al., 2000; R. A. L. Sia et al., 2003). We find evidence that translesion synthesis initiated by the *REV1* gene during DNA replication is responsible for the nuclear mutation rate difference between haploid and diploid *S. cerevisiae*; upon deleting *REV1*, we observed the single-nucleotide mutation rate decrease by 58.7% in haploids and 33.5% in diploids, such that the SNM rates of haploid and diploid *rev1*Δ yeast were statistically indistinguishable (Fig. 1A).

Among our *rev1*Δ lines, we found that on average, mutations occurred at earlier-replicating sites in haploids and later-replicating sites in diploids compared to wild type lines (Fig. 1C). However, we failed to detect a significant interaction effect, likely due to reduced mutation rates and subsequently smaller samples of mutations among our *rev1*Δ treatments. This trend is consistent with previous research by Sharp et al. (2018), who found that the ploidy difference in SNM rates depended on replication timing, with an elevated mutation rate specifically in late-replicating regions in haploids. The Rev1 polymerase has been found to be highly expressed during G_2_/M phase of the cell cycle in haploids, after most DNA has been replicated (Waters & Walker, 2006), and there is evidence for a mutagenic effect of Rev1 in late-replicating genomic regions in haploids (Lang & Murray, 2011). Our data suggest that the timing of Rev1 activity may depend on ploidy.

We found evidence that deleting *REV1* caused a significant decline in C to G transversions in both haploids and diploids (Fig. 2A). This is consistent with previous studies that have found the Rev1 polymerase to be a major driver of C to G SNMs in multiple taxa (Kano et al., 2012; Larimer et al., 1989; Sasatani et al., 2020). Our data, which found depressed rates of this mutation type in both cell types in the absence of *REV1*, suggest that haploids and diploids both employ TLS as a strategy to complete DNA replication before reaching the G2/M boundary of the cell cycle (Foiani et al., 2000; Putnam et al., 2009). Taken with the interaction effect between ploidy and *REV1* genotype on the nuclear SNM rate that we found, our experiment suggests that *REV1* confers a stronger mutator phenotype in haploids than in diploids. Both haploid and diploid cells are known to utilize an alternative repair pathway, template switching (TS), through homologous recombination with the sister chromatid to overcome errors during DNA replication (Smith et al., 2007; Štafa et al., 2014), and specific genes responsible for sister chromatid recombination have been identified (Mozlin et al., 2008; Muñoz-Galván et al., 2012). It is possible, however, that haploids express *REV1* earlier in the cell cycle compared to diploids, or that the threshold for the employment of TLS instead of TS is lower in haploids. Future work could elucidate the pattern of transcription of this gene as well as other DNA repair genes throughout the cell cycle to further understand what causes TLS to be more mutagenic in the less common ploidy type. While data exist on temporal gene expression patterns in haploid *S. cerevisiae* (Kingsman et al., 1985; Mitchell, 1994), comparisons of haploid and diploid gene expression are sparse.

Following the deletion of *REV1*, we observed an increase in the growth rate of haploids, but not diploids, when grown in YPAD (Fig. 3A). Taken with the decreased nuclear SNM rate of *rev1*Δ treatments compared to their wild-type counterparts, this is a somewhat surprising result in haploids; the Rev1 polymerase is known to assist cells in the completion of DNA replication and therefore cell division (Lawrence, 2002), so we expected cells with functional *REV1* gene function to grow faster in both ploidy types. Given that *rev1*Δ MA lines (i) harbored fewer nuclear mutations and (ii) grew at a similar or faster rate, we wanted to better understand what benefit this gene might provide. We tested whether the *rev1*Δ strains might exhibit reduced fitness with respect to UV resistance (Fig. 3B) or in direct competition with their *REV1*+ counterparts in the presence of the oxidative stressor t-BOOH (Fig. 3C). We found that both the haploid and the diploid *rev1*Δ lines had higher resistance to UV exposure; this is partly consistent with evidence that *REV1* is not required for replication past T – T (6 – 4) UV photoproducts (Nelson et al., 2000; Otsuka et al., 2005). We also found that *rev1*Δ strains showed a competitive advantage in the presence of t-BOOH. To our knowledge, there is no prior evidence that tolerance of mutagenic stressors should improve without a functional copy of *REV1*. In summary, these data did not clarify the fitness benefit of maintaining a functional copy of *REV1*.

Given that the loss of *REV1* would lead to a decreased nuclear mutation rate, without any apparent cost to growth or mutagen resistance, we should ask why this gene has been maintained by selection. The answer may lie in the role of *REV1* in mitochondrial genome maintenance. We found that *rev1*Δ lines of both cell types were significantly more likely to become petite; while all 100 *REV1*+ lines successfully grew on non-fermentable media following MA, five haploid and seven diploid *rev1*Δ lines did not (Fig. 1D), indicating a block in the aerobic respiratory chain pathway (Day, 2013). Further analysis of the mitochondrial sequence data showed that 11 of the petite lines had lost their mitochondrial genomes entirely, while one had a loss-of-function mutation at the *COX1* gene, which is known to confer a petite phenotype (Lemaire et al., 1998; Meunier et al., 1993). Additionally, we found a significant increase in the mitochondrial point mutation rate among our *rev1*Δ lines (Fig. 1B), providing further evidence that *REV1* contributes to mitochondrial stability, perhaps at the expense of nuclear genome replication fidelity. This contradicts previous research by Zhang et al. (2006), who used the *MtArg8* reversion assay to demonstrate a reduced point mutation rate in the mitochondria of Rev1-deficient haploid yeast cells. However, our findings are consistent with other research on the topic, which suggests that the Rev1 polymerase contributes to proper mitochondrial function and mitochondrial genome stability (In Het Panhuis et al., 2022; Rasmussen, 2003). Another study found that Rev1-deficient mouse cells displayed signatures of mitochondrial dysfunction according to measurements of electron transport chain activity and oxidative phosphorylation (Fakouri et al., 2017).

Taken as a whole, the experimental data from our study contribute to a body of work that elucidates the evolutionary tradeoffs associated with the *REV1* gene. We observed significantly stronger maintenance of mitochondrial genomes in MA lines with a functional copy of *REV1*, at the apparent cost of reduced nuclear genome replication fidelity and decreased resistance to mutagenic stress. These tradeoffs were consistent across haploids and diploids, but we observed an additional fitness cost of *REV1* with a ploidy-specific effect: *REV1*+ haploids grew significantly more slowly than their *rev1*Δ counterparts in YPAD, a relevant fitness cost in yeast (Miller et al., 2022). The fact that this effect was only observed in the rare cell type is consistent with the idea that haploid growth has been subjected to limited selection in the history of this predominantly diploid organism.

The drift-barrier hypothesis posits that purifying selection acts inefficiently against mutator alleles with small fitness effects relative to N_e_ (Lynch et al., 2016). An extension of this idea is that purifying selection acts inefficiently against mutator alleles that only affect a rare cell type (Bao et al., 2025; Sharp et al., 2018). Our results suggest that the *REV1* gene has particularly detrimental effects on haploid *S. cerevisiae*, increasing mutation rate and reducing growth rate, and that this effect persists because selection on this rare cell type has historically been less effective. The *S. cerevisiae* genome could include other alleles that are deleterious to haploids, but selectively neutral or beneficial in diploids. Given that a single gene’s function appears to cause a difference in a fundamental biological phenomenon depending on cell type, we recommend that ploidy be given greater weight as a variable in future genetic studies, particularly in systems such as *S. cerevisiae*, where cells can readily be grown in either form.

---

**Supplemental figure 1.** Fluorescence peaks of cells fixed with SYTOX Green, a fluorochrome used to stain and quantify DNA, showed that spontaneous ploidy change was rare. By comparing data from each MA line with haploid (red) and diploid (blue) controls, we found two haploid lines that became diploid over the course of MA.

## Data availability

Sequence data are available from the National Center for Biotechnology Information Sequence Read Archive (accession no. PRJNA1337143). R scripts used to filter sequence data, analyze results, and generate figures are available on figshare (DOI 10.6084/m9.figshare.31079359). Supplementary table 1 is also available on figshare (DOI 10.6084/m9.figshare.31100323). Additional data are available as supplementary information.

## Acknowledgements

We thank L. Hoffman, B. Lieser, and L. Margolin for assistance with pouring plates and conducting growth rate measurements, and K. Guerra and C. Oguejiofor for developing protocols for mutagen assays. This work was supported by the National Institute of General Medical Sciences of the National Institutes of Health under award number R35GM154954 to NPS, and the American Genetic Association under the Evolutionary, Ecological, and Conservation Research Award to JF. The authors used the University of Wisconsin-Madison Biotechnology Center’s DNA Sequencing Facility (RRID:SCR_017759) and the University of Wisconsin Carbone Cancer Center Flow Cytometry Laboratory, supported by P30 CA014520 from the National Cancer Institute of the National Institutes of Health.

